# Microfluidic Capture-Enrichment of individualized microbes from urban wastewater reveals a hidden reservoir of potential pre-emergent pathogens

**DOI:** 10.64898/2026.09.21.753334

**Authors:** Siyi Zhou, Seokju Seo, Kathryn Rhatigan, Marga Lee, Alice Liu, Alyse Tran, Fernanda Vaca, Lois Armendariz, Todd Treangen, Natalia V. Kirienko, Lauren B. Stadler, Yousif Shamoo

**Affiliations:** Department of Civil and Environmental Engineering, Rice University, 6100 Main Street, Houston, TX, 77005, United States of America; Department of BioSciences, Rice University, 6100 Main Street, Houston, TX, 77005, United States of America; Department of Computer Science, Rice University, 6100 Main Street, Houston, TX, 77005, United States of America

**Keywords:** Environmental surveillance, phenotyping, whole genome sequencing, emerging pathogens, microfluidics, human serum resistance, innate immune evasion

## Abstract

Current environmental biosurveillance infrastructure excels at tracking known biological threats but is limited in its ability to proactively identify potential pre-emergent pathogens (PEPs). To address this gap, we developed a microfluidics-based Capture-Enrichment pipeline that isolates individual environmental cells via microdroplet encapsulation, preserving community biodiversity while enabling iterative selection against human sera. When applied to urban wastewater, this approach successfully recovered pathogens, commensals, and PEPs that fell below the detection limits of conventional amplicon sequencing. Genomic analysis revealed that wastewater-derived ESKAPE pathogens cluster closely with clinical isolates, showing that the pipeline captures clinically relevant threats. Furthermore, isolated PEPs exhibited known and emerging phenotypes associated with early-stage pathogenesis and antimicrobial resistance. These findings demonstrate that targeting PEPs can fundamentally expand the scope and quality of biosurveillance by providing a scalable strategy to identify, characterize, and close important gaps in our understanding of emerging microbial pathogens.

## Introduction

As a consequence of global climate change and the continuing expansion of the human population, the number of emerging pathogens is predicted to expand^1,2^. A challenge for public health is to understand and identify these unknown emerging pathogens before they cause clinical harm. While the definition of a pathogen is fairly clear, denoting any organism or infectious agent that causes disease, our practical ability to classify environmental microbes as pathogens is not^3^. This biological ambiguity underscores a fundamental challenge in pathogen classification and suggests that expanded phenotypic characterization of environmental strains could dramatically improve sequence-based biosurveillance.

Environmental surveillance of infectious diseases, such as wastewater monitoring, relies on genomic methods to target or flag organisms that carry known virulence factors, resistance genes, or homology to characterized pathogens^4,5^. This approach can only flag what has already been annotated as dangerous, and cannot determine whether a novel organism is capable of causing disease, because pathogenic potential is a functional property that, at present, cannot be predicted from sequence alone^6–8,9,10^. Strains with host-infection capacity can therefore circulate undetected until they cause substantial disease, leaving surveillance reactive rather than preemptive^11^. Closing this gap requires a scalable, phenotypic method that can directly test environmental microbes for pathogenic potential, independent of genomic novelty or prior clinical recognition.

In this study, we use a phenotypic approach to identify strains as "Potential Pre-Emergent Pathogens" (PEPs) based on a primary physiological marker: the capacity to grow in human serum. We hypothesize that the capacity to overcome the powerful antimicrobial properties of human serum serves as a critical, early indicator of pathogenic potential, regardless of an organism’s genomic novelty or current clinical recognition^3,12^. It functions as a non-permissive environment for microbes unequipped to survive within it. As a primary component of the innate immune response, serum eliminates many environmental microbes through a formidable immunological arsenal that includes complement, antibodies, antimicrobial peptides, and nutritional immunity^13^.

To harness this evolutionary selection filter for PEP discovery, we developed "Capture-Enrich," a high-throughput platform driven by microfluidic encapsulation. Capture-Enrich uses the growth of individualized cells in human serum as both an initial indicator of concern for a PEP strain and a physical means to enrich rare microbes by down-selecting against abundant, non-pathogenic environmental microbiota. Our results reveal a substantial reservoir of PEPs within the environment and present a scalable, strain-agnostic framework to capture, analyze, and functionally validate uncurated PEP strains. We show that wastewater can harbor an under-surveilled, clinically adjacent reservoir of opportunistic lineages possessing active host-interaction phenotypes and multi-drug resistance. By exploring the structural gaps between these observed functional traits and existing genomic annotations, we can enrich reference databases and provide the biological context necessary to advance sequence-based biosurveillance.

## Results

### Capture-Enrich platform selects for microbes that can flourish under selection by human serum

Capture-Enrich is a high-throughput platform that uses microfluidic encapsulation (Fig. 1) to uncover latent biodiversity by isolating and enriching individual strains. Spatial confinement within microdroplets protects slow-growing species from being outcompeted by faster-growing counterparts, limiting cellular expansion to the carrying capacity of each independent droplet^14,15^. We collected samples from two geographically distinct locations in Houston selected for their potentially divergent environmental microbiomes. Site A is a regional wastewater treatment plant (WWTP) that receives effluent from both residential areas and the Texas Medical Center (TMC). As one of the largest medical complexes globally, with over 10 million patient interactions annually, Site A may represent a unique catchment for emerging pathogens. Site B, in contrast, captures a standard, mixed-residential profile.

**Fig. 1.**
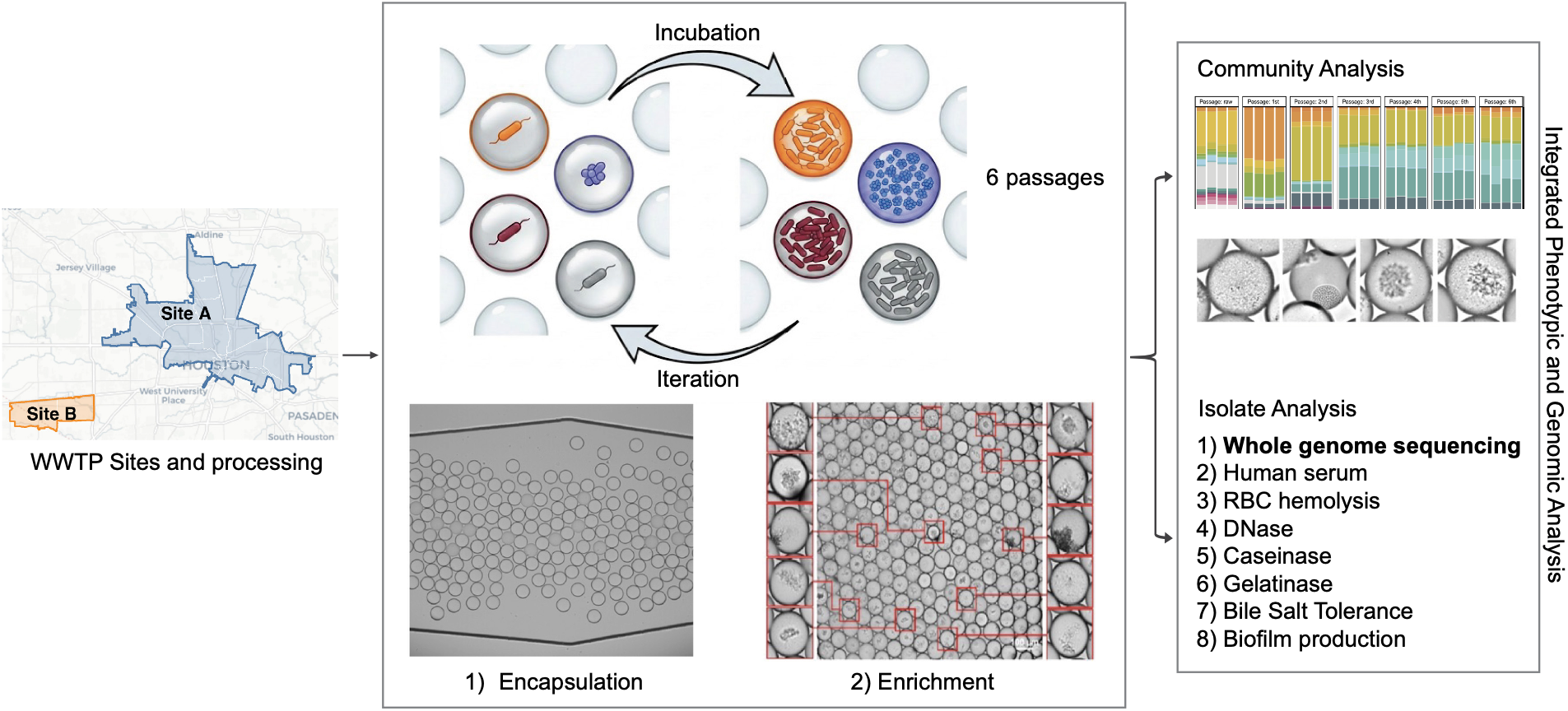
The Capture-Enrich platform. Wastewater from two Houston-area treatment plants (Site A, blue, receiving Texas Medical Center effluent; Site B, orange, mixed-residential) was encapsulated in ∼90 µm human-serum microdroplets at 0.3–1.0 cells per droplet and serially passaged against serum for six iterations over 18 days (four replicate populations per site; scale bar, 100 µm). Decapsulated populations were tracked across passages (microbial community analysis), and 161 recovered isolates were characterized by whole-genome sequencing and phenotypic assays (isolate analysis). To evaluate host-pathogen interactions, a subset of nine candidate PEPs was subsequently screened for *in vivo* virulence using *Caenorhabditis elegans* pathogenesis assays.

We used growth in human serum as the enrichment condition. Cell was encapsulated directly from wastewater as a microdroplet emulsion with an average of 0.3-1 cells/microdroplet in human serum. Under these conditions, each 1 mL emulsion of 90 µm microdroplets captured approximately 566,800 to 965,800 individualized cells (Table S1). To assess population dynamics within our system over an 18-day period, we passaged four replicate populations from each site as 1 mL emulsions for six iterations (Fig. 1). Following each 3-day incubation period at 37°C, we decapsulated the microdroplets, diluted the recovered cells, and re-encapsulated them with fresh serum for the subsequent round of enrichment. After each passage, we measured the optical density of each decapsulated population. The average growth per iteration ranged from 3×10^7^ and 2×10^9^ cells/ml, consistent with robust growth of the microbial populations within microfluidic droplets.

### Serial serum passaging exposes a hidden reservoir of serum-prolific wastewater bacteria

We used 16S rRNA gene amplicon sequencing to characterize the microbial communities from Sites A and B. Serum exposure produced an immediate diversity bottleneck at both sites: After one passage, observed ASV richness decreased from 2,805 ± 285 at Site A and 1,528 ± 61 at Site B in raw wastewater to 489 ± 62 and 451 ± 95, respectively, corresponding to a 70–83% reduction in richness (Friedman tests, p = 0.005 (Site A) and p = 0.003 (Site B); Fig. 2a; Fig. S1a, b). Passage was the dominant driver of community restructuring, explaining 76.2% of compositional variation in Bray–Curtis ordination, whereas site explained 9.1% (PERMANOVA, marginal tests, p < 0.001; Fig. 2b), whereas site accounted for 9.1%. Both sites followed a shared serum-selection trajectory, but the communities did not converge. Instead, between-site separation increased by the final passage, indicating that serum imposed a common functional filter while allowing distinct local microbial communities to emerge (Fig. S1c, d).

**Fig. 2.**
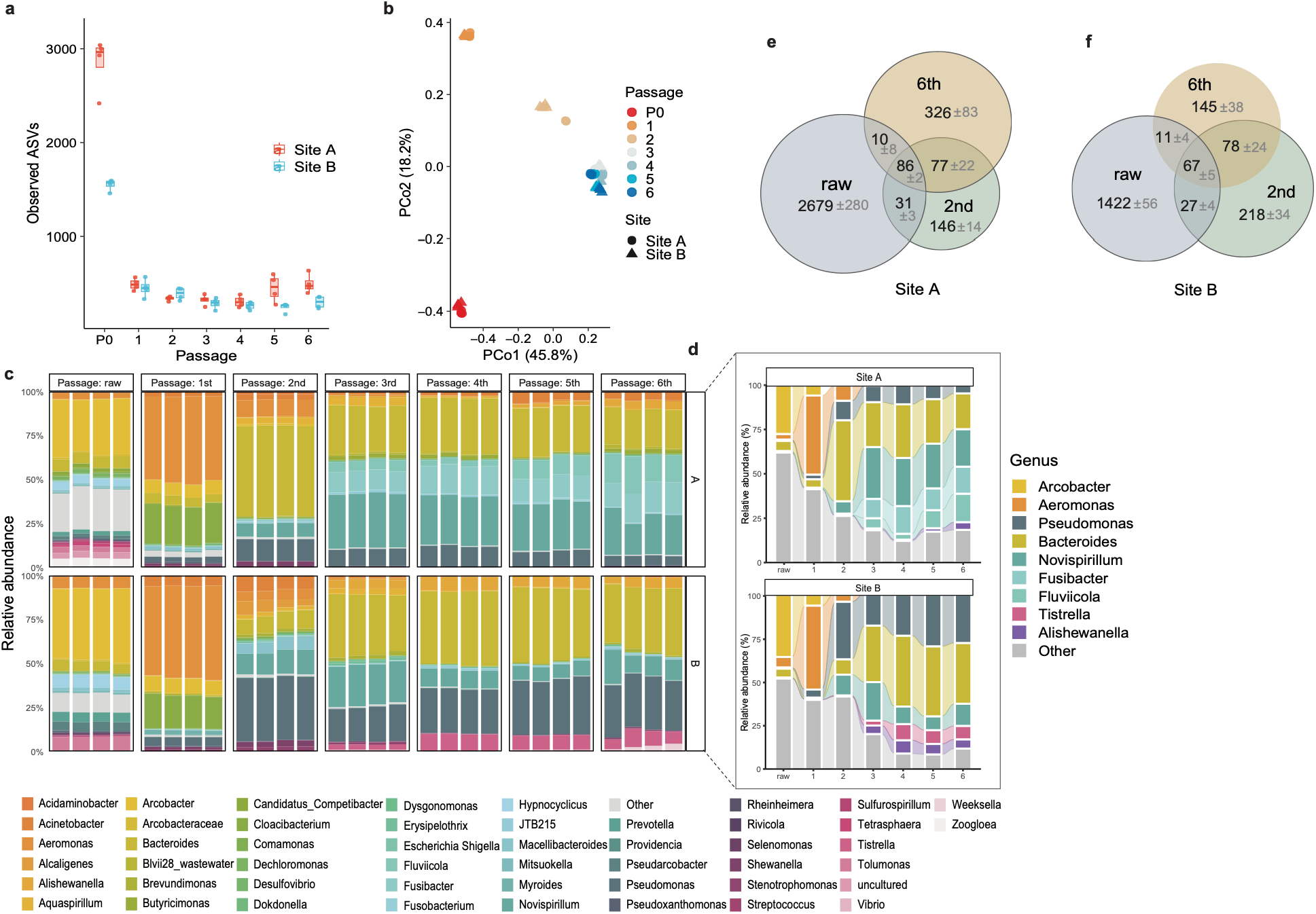
Serial serum passaging restructures wastewater microbial communities. Each site was sampled once and the sample divided into four aliquots, each founding one parallel enrichment lineage carried through six serial passages; all panels show these four lineages per site. (a) Observed ASV richness across raw wastewater (P0) and six serial serum passages (P1 to P6) at two WWTP sites (Site A, red; Site B, blue). Points show individual lineages; boxplots show median and interquartile range. Because the seven timepoints within a lineage are repeated measures on one propagated population, the effect of passage was tested by Friedman tests blocked on lineage. (b) Principal coordinates analysis of Bray-Curtis dissimilarity of ASV-level relative abundances. Points are colored by passage and shaped by site. PERMANOVA with marginal terms and 9,999 permutations, restricted within lineage: passage R² = 0.76, p < 0.001. Site accounted for R² = 0.09; because each site is represented by a single wastewater sample, this is reported as an effect size without an associated p value. (c) Genus-level relative abundance in each lineage. Genera were ranked by overall relative abundance across all samples; the 50 most abundant are shown individually and all others grouped as Other. (d) Passage-level trends for selected genera from c, shown separately for each site as the mean of four lineages. Ribbons connect the same genus across passages and all other genera are grouped as Other. Genera were selected for change across passages rather than for overall abundance: *Novispirillum, Fusibacter, Fluviicola, Tistrella* and *Alishewanella* were each present below 1% in raw wastewater. (e, f) Overlap of ASVs detected in raw wastewater, passage 2 and passage 6 at Site A (e) and Site B (f), each lineage compared against its own raw aliquot. Values are mean ASV counts ± s.d. across the four lineages.

Taxonomic succession revealed that serum passaging did not simply enrich the most abundant raw-wastewater taxa. *Arcobacter* dominated raw wastewater but declined to ≤1% at both sites by passage 2. *Aeromonas* bloomed transiently after one passage, reaching 44.2 ± 1.0% (Site A) and 47.4 ± 0.9% (Site B), but collapsed below 9% by passage 2. From passage 2 onward, *Bacteroides* expanded from under 6% in raw wastewater to 20.0 ± 0.3% (Site A) and 34.3 ± 2.2% (Site B) by passage 6 (Fig. 2c, d; Table S2). Late-passage communities also retained site-specific signatures: *Pseudomonas* was a major component of Site B (28.5%) but minor at Site A (5.1%), and *Acinetobacter* was enriched at Site A (4.4% vs 0.8%) (Fig. 2c, d).Taken together, these results show that serum passaging drove reproducible taxonomic succession while preserving site-specific community signatures.

Serum passaging also recovered lineages that fell below the detection limit of direct amplicon sequencing of raw wastewater (mean 249,235 ± 44,344 quality-filtered reads per sample; Fig. S1a). Raw wastewater, passage 2, and passage 6 shared only a limited number of ASVs: 86 ± 2 ASVs at Site A and 67 ± 5 at Site B (Fig. 2e, f), out of the ∼2,800 (Site A) and ∼1,530 (Site B) ASVs present in raw wastewater. By passage 6, only 25.1 ± 3.1% of Site A and 36.3 ± 5.2% of Site B ASVs detected were present in their original raw wastewater; the remaining 373 ± 90 (Site A) and 194 ± 55 (Site B) ASVs were present at passage 6 and not in the corresponding raw wastewater sample (Table S3). Their recovery under enrichment suggests they were present in raw wastewater at abundances below our detection limit. Across the passage series, enrichment recovered taxa from 27 and 31 genera at Sites A and B that were undetected in raw wastewater (Table S4). Enrichment also expanded 21 genera that were initially at frequencies <1% in raw wastewater (Fig. S2). These findings show that serum passaging exposes rare wastewater lineages that direct sequencing of raw wastewater does not detect, and that a subset of these persist under sustained serum pressure.

### Serum-selected isolates are under-surveilled but related to clinically sampled lineages

To evaluate and compare the strains enriched from wastewater samples by Capture-Enrich to known isolates, whole genome sequencing was performed on 161 isolates recovered from passage 2 across both sites. Passage 2 was selected because a transient *Aeromonas* bloom had collapsed by this point, from 44.2 ± 1.0% and 47.4 ± 0.9% at passage 1 to 8.5% (Site A) and 5.1% (Site B) (Fig. 2c, d; Table S2). Whole-genome sequencing identified 31 species across 17 genera, with *Pseudomonas aeruginosa* (n = 32) and *Alcaligenes phenolicus* (n = 26) as the largest species groups (Fig. 3a; Table S5). The collection contained both repeated local expansions and diverse singletons: 71 isolates (44%) fell into 21 clonal groups defined by ≥99.9% ANI and ≤100 core SNPs, and no clonal group spanned both sites (Table S6). The collection also included candidate novel taxa, including a four-isolate *Alishewanella* lineage with 85–86% ANI to the nearest described species and a *Pseudomonas* isolate with ∼88% ANI to its nearest described relatives. Overall, 105 of 161 isolates either carried novel sequence types (STs) or belonged to genera without an established multilocus sequence typing (MLST) scheme, placing much of the serum-prolific reservoir outside standard sequence-type surveillance frameworks (Table S5).

**Fig. 3.**
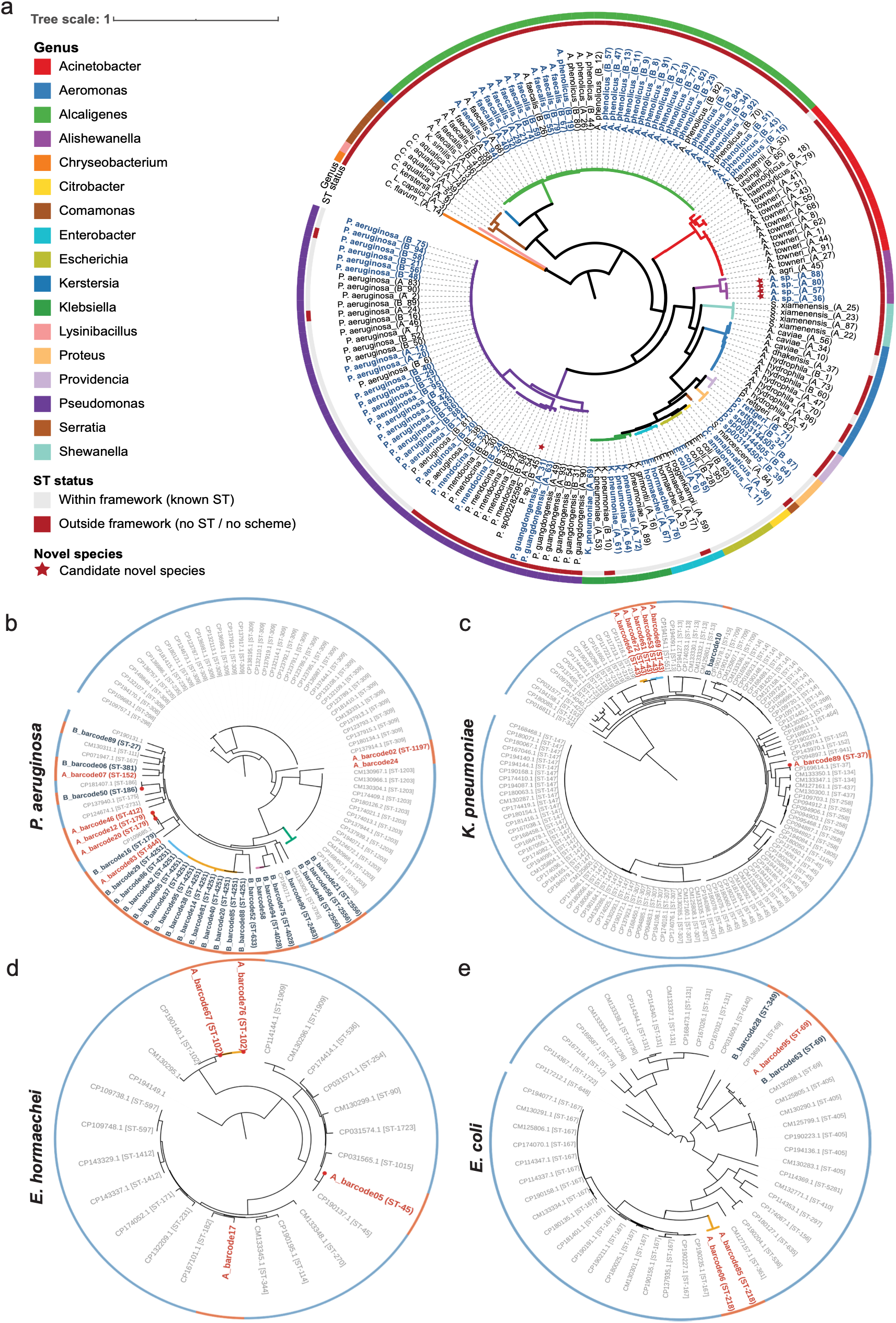
Serum-enriched isolates are under-surveilled and clinically adjacent. (a) Phylogenomic tree of the 161 serum-enriched wastewater isolates, inferred with GTDB-Tk from the concatenated bac120 marker-gene alignment and visualized in iTOL. Tip labels give the species assignment and isolate origin (A = Site A; B = Site B). Branch color and the outer ring denote genus (legend). Tip-label color indicates clonal-group membership: isolates belonging to a clonal group (≥99.9% ANI and ≤100 core SNPs) are shown in blue, singletons in black. The inner ring indicates sequence-type surveillance status: isolates within current frameworks (known ST; white) versus outside them (novel/unassigned ST or a genus with no MLST scheme; red); 105 of 161 isolates (65%) fall outside. Red stars mark the two candidate novel species (<95% ANI to the nearest described species in GTDB): a novel *Alishewanella* recovered four times independently at Site A (85–86% ANI to *A. agri* / *A. aestuarii*) and a single novel *Pseudomonas* at Site B (∼88% ANI to *E. oleovorans* / *E. mendocina*). (b) - (e) Wastewater isolates embed within clinical isolate phylogenies for the four most abundant ESKAPE-adjacent species. Maximum-likelihood phylogenies of P. aeruginosa (32 wastewater + 59 clinical), *K. pneumoniae* (7 + 48), *E. hormaechei* (4 + 22), and *E. coli* (5 + 50) genomes. Outer ring indicates source: wastewater isolates (red) and CDC HAI-Seq clinical assemblies (blue; BioProject PRJNA288601). Branch colors denote wastewater clonal group membership. Per-species core-gene alignments were built with Panaroo (strict mode, 0.95 core threshold) and trees inferred with RAxML (GTRGAMMA) and visualized in iTOL. Wastewater isolates interleave with clinical references rather than forming distinct environmental clades; 44 of 48 isolates (92%) share ≥99.0% ANI with their nearest clinical isolate genome. Five wastewater isolates shared sequence-type assignments with their nearest CDC HAI-Seq clinical isolate genomes; pairwise distances ranged from 157 to 310 SNPs across the shared core genome.

We next tested whether Capture-Enrich recovered lineages were related to clinically sampled bacteria. For the most abundant ESKAPE-adjacent species (*P.* aeruginosa, *K. pneumoniae*, *E. hormaechei* and *E. coli*), wastewater isolates were embedded into reference phylogenies built from CDC HAI-Seq clinical assemblies (Table S7). Wastewater isolates interleaved with clinical isolate genomes rather than forming environmental-only clades, and 44 of 48 wastewater isolates from these selected species shared ≥99.0% ANI with their nearest clinical isolate genome (Fig. 3b–e; Table S5). Several wastewater isolates matched STs observed among CDC HAI-Seq clinical isolate genomes, including *E. hormaechei* ST45/ST102, *K. pneumoniae* ST37 and *P. aeruginosa* ST186 (Table S7). For these isolates, the nearest clinical isolate genomes differed by 157–310 SNPs across the shared core genome, suggesting close but non-identical relatedness. Two *E. coli* ST69 isolates were more dissimilar to the closest clinical isolate genomes (CM130288.1) at 3,380 and 1,845 core SNPs, highlighting broader within-ST diversity in this globally disseminated extraintestinal pathogenic *E. coli* (ExPEC) lineage. Notably, clinical adjacency did not always coincide with canonical markers of concern: all seven *K. pneumoniae* isolates lacked Kleborate hypervirulence and high-priority resistance signatures and their capsule types (K12, K30, K45, K103) fell outside the loci associated with hypervirulent (K1, K2, K20) or carbapenem-resistant high-risk (K15, K17, K28) lineages (Table S5)^16^. Serum enrichment, therefore, recovered wastewater lineages that were closely related to clinically sampled bacteria by genome-wide comparisons but would not necessarily be prioritized by current marker-based surveillance.

### Serum-enriched isolates include both tissue-damaging strains and strong serum growers with low damage activity

To define phenotypes that correlated with strong growth in serum, we assayed 161 aerotolerant isolates for seven *in vitro* traits spanning tissue damage, colonization, and host-environment persistence: hemolysis, gelatinase, caseinase, DNase activity, biofilm formation, bile-salt tolerance, and growth in 100% human serum (Fig. 4a; Table S8). The isolates did not follow a single virulence gradient. Bile-salt tolerance was the most common phenotype (129 of 161 isolates). Tissue-damage phenotypes were less frequent: 55 isolates were gelatinase-positive, 45 were caseinase-positive, 44 were hemolytic and 23 were DNase-positive. We also performed serum-growth assays for each isolate. While all isolates grew well enough in serum to proliferate over 6-passages in microdroplets, variations in serum cloudiness for the microplate-based assay limited us to only include ODs >0.1 after baseline subtraction (Fig. S3; Table S8). We summarized non-serum phenotypes using a damage-weighted phenotype index (DPI) calculated from binary phenotype calls, with hemolysis, gelatinase, caseinase, and DNase activity weighted more heavily than biofilm formation and bile-salt tolerance (Methods; Fig. S4). DPI correlated strongly with serum growth (Fig. 4b; Spearman’s ρ = 0.64, P < 0.001), but 13 isolates grew well in serum despite a low DPI (Fig. 4c). To investigate drivers of damage phenotypes, we evaluated the taxonomies of the high- and low-DPI enriched bacteria. *P. aeruginosa* (n = 32) showed a high-damage profile, with uniform hemolysis and gelatinase activity (32/32) and frequent caseinase activity, biofilm formation, and bile-salt tolerance (29–30/32; mean DPI 7.7), whereas *A. hydrophila* (n = 7) showed broad tissue-damage activity (Fig. 4d). Principal-component analysis of the phenotype profile separated isolates largely by taxonomy along PC1 (50.6%), which increased with DPI, with *Pseudomonas* and *Aeromonas* at the high-DPI end (Fig. 4e). PC2, (17.1%) was independent of DPI and was defined by bile-salt tolerance and DNase activity, indicating an axis of phenotypic variation that the DPI does not capture (Table S9). Species explained most of the phenotype variance (PERMANOVA R² = 0.68) while sampling site explained essentially none (R² ≈ 0.00, n.s.). High-DPI isolates were dominated by recognized opportunistic pathogens such as *P. aeruginosa*, *A. hydrophila*, and *K. pneumoniae*, whereas low-DPI isolates were taxonomically more heterogeneous and included the serum-prolific strains described above. Hemolysis, gelatinase, and caseinase showed the strongest positive pairwise correlations (Fig. 4f; Spearman ρ = 0.79–0.84), whereas DNase, biofilm formation and bile-salt tolerance showed weaker or more trait-specific relationships. Although all isolates were recovered following the same serum-droplet selection, downstream phenotyping resolved two coexisting groups: recognized opportunistic pathogens with coordinated tissue-damage activity, and a low-DPI serum-prolific group lacking these phenotypes. Serum growth thus defined traits separable from canonical tissue damage.

**Fig. 4.**
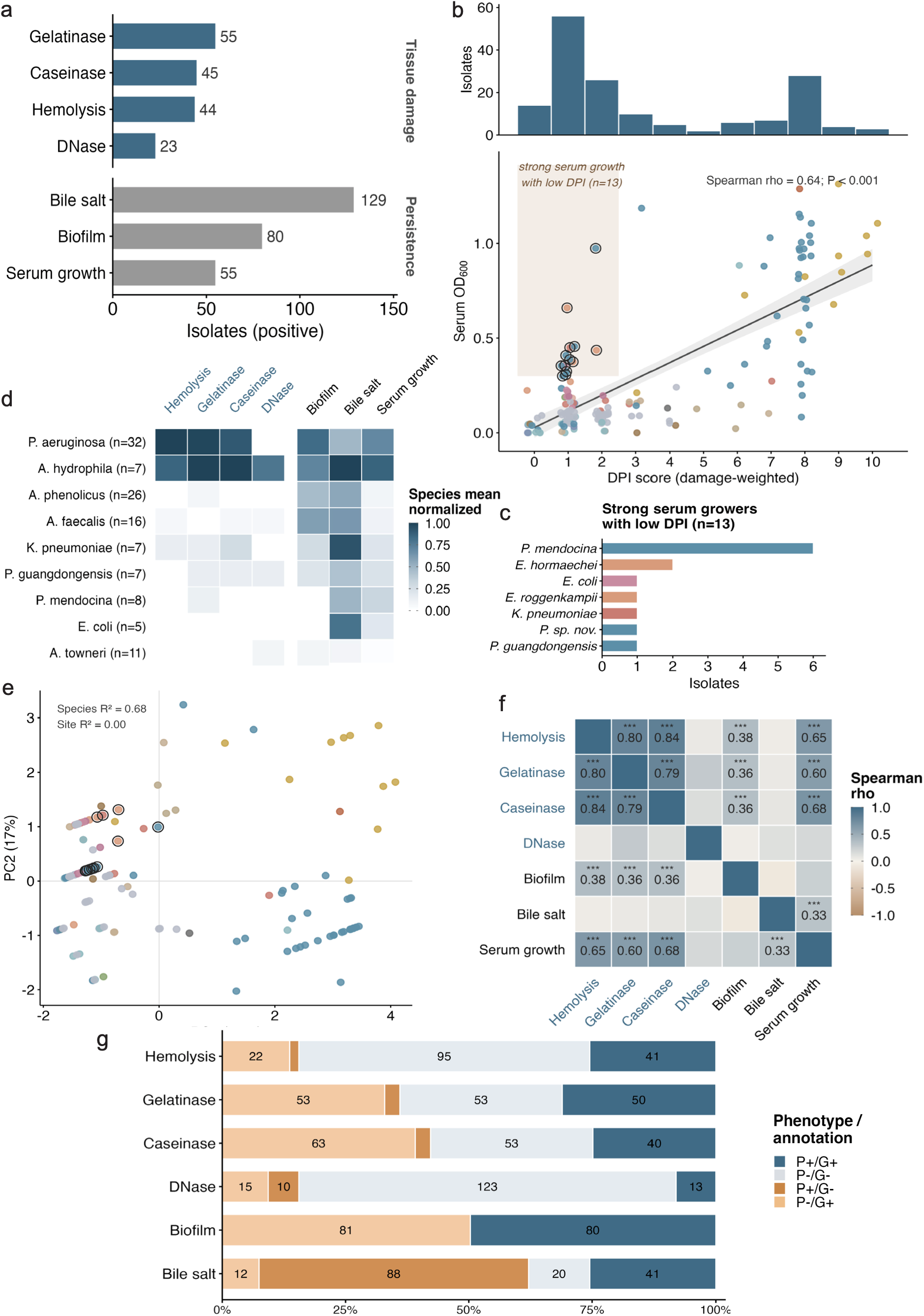
Serum-enriched isolates resolve into damage-associated and low-damage serum-prolific states. (a) Number of isolates positive for each of the seven assays, grouped into tissue-damage and persistence traits; tissue-damage bars are shown in blue. (b) Damage phenotype index (DPI) across the collection. Top, distribution of DPI scores. The distribution appeared bimodal, with a large mode at DPI 0 to 2, a second mode at DPI 8, and a trough at DPI 5 separating them. Bottom, relationship between DPI and serum growth; points are colored by genus, the shaded region and ringed points mark isolates combining strong serum growth with minimal damage activity (DPI ≤ 2, n = 13), and the Spearman correlation between DPI and serum growth is indicated. (c) Taxonomic composition of the low-DPI serum-prolific isolates. (d) Species-level mean normalized phenotype profiles for the nine species represented by at least five isolates, across the seven assays grouped into tissue-damage and persistence traits (columns); tissue-damage assay labels are shown in blue. (e) Principal-component analysis of the phenotype profile, with PC1 oriented to increase with DPI. Points are colored by genus and the 13 low-DPI serum-prolific isolates are ringed. Serum growth loaded on PC1 (0.45) alongside caseinase, hemolysis and gelatinase (0.48 to 0.49), whereas PC2 was defined by bile-salt tolerance (0.68) and DNase activity (0.55) opposed to biofilm formation (−0.44). Marginal PERMANOVA R² for species and site is annotated. (f) Spearman correlations among the seven assays. Hemolysis, gelatinase and caseinase were the most strongly correlated (ρ = 0.79 to 0.84); tissue-damage assay labels are shown in blue. (g) Concordance between each observed phenotype and the presence of a curated trait-linked gene. Bars give the fraction of isolates in each category: phenotype-positive and gene-positive (P+/G+), phenotype-negative and gene-negative (P−/G−), phenotype-positive and gene-negative (P+/G−), and phenotype-negative and gene-positive (P−/G+). Numbers are isolate counts. The genus color key applies to panels b, c, and e.

To test how reliably sequence data predicted these functional traits, we compared each isolate’s phenotype to the presence of characterized candidate genes in its genome (Fig. 4g; Table S12, Table S13). Genotype reliably predicted the four tissue-damage phenotypes (hemolysis, gelatinase, caseinase, and DNase; all p < 1×10⁻⁴) but failed for persistence and serum-survival traits; none of the 55 strong serum growers, including 44 ESKAPE-adjacent genera, carried a curated serum-survival or complement-evasion marker gene. This discordance shows that curated gene sets capture overt tissue-damage traits but miss the serum-survival phenotypes that define this collection, reinforcing that marker-gene screening alone cannot substitue for functional testing.

### Functional screening of wastewater isolates demonstrates pathogenic potential

Nine strains were selected for deeper phenotypic characterization that spanned the DPI (0-7) and are poorly characterized in the literature: *Comamonas kerstersii, P. guangdongensis, Citrobacter amalonaticus, Acinetobacter towneri, Shewanella xiamenensis, Lysinibacillus capsici, Chryseobacterium flavum, E. roggenkampii* and *Alishewanella sp. nov..* To evaluate their pathogenic potential, we used a *C. elegans* infection model, exposing *C. elegans glp-4(bn2)* worms to each isolate in an agar-based assay, analogous to a Slow Killing assay developed for *P. aeruginosa*. All tested isolates significantly reduced host survival relative to *E. coli* (Fig. 5a). Median survival time (LT50) varied across strains, reflecting a range of pathogenic potential (Fig. 5b), but consistent with potential pathogenicity.

**Fig. 5.**
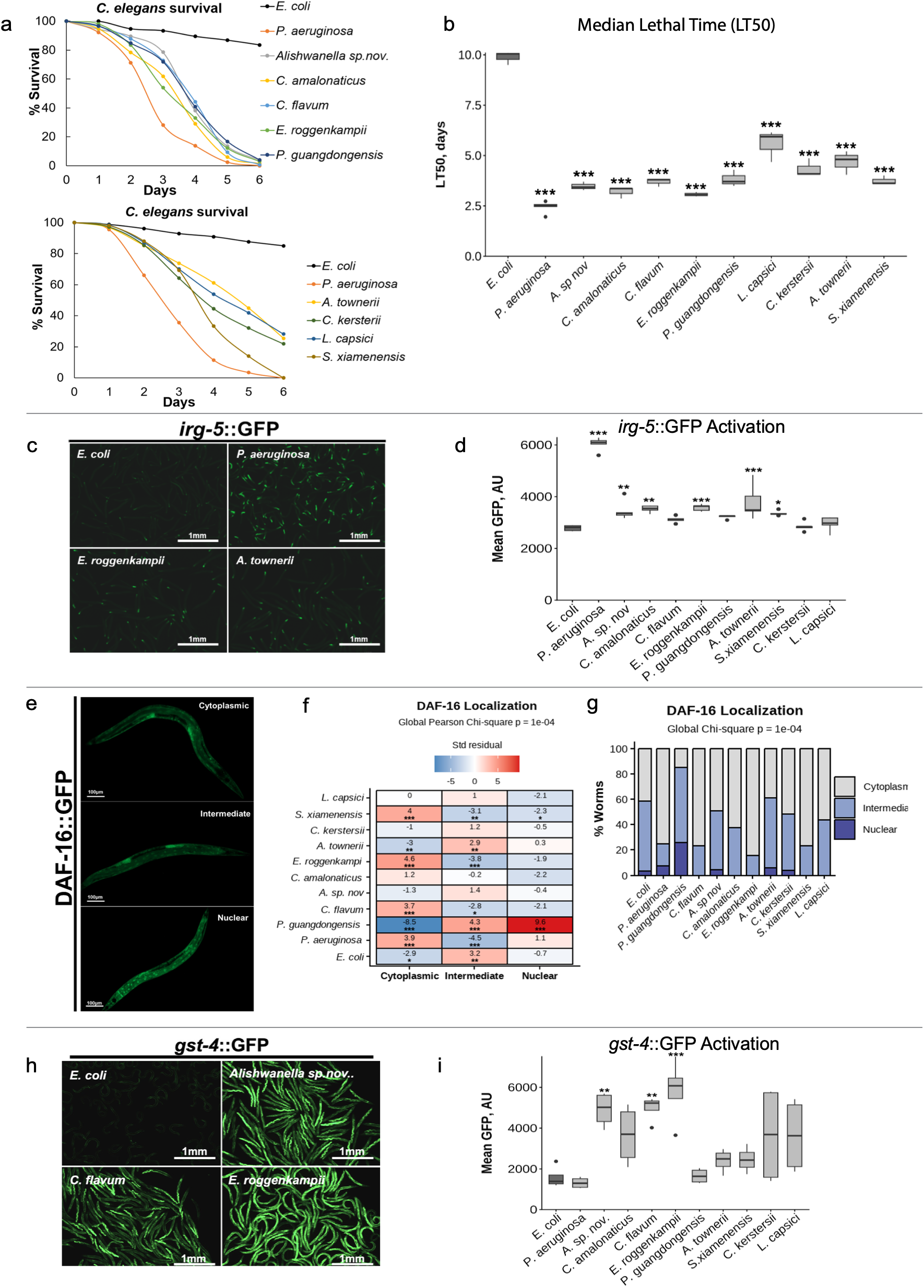
Wastewater isolates affect C. elegans survival and trigger host-defense pathways. (a) Survival of *C. elegans* exposed to different wastewater isolates across 6 days. (b) LT50 (median time to 50% mortality) were calculated from the corresponding survival curves. (c) Fluorescent images of *C. elegans* carrying *irg-5p*::GFP exposed to *E. coli*, *P. aeruginosa*, *E. roggenkampii*, or *A. townerii*. (d) Quantified mean fluorescence intensity of *irg-5p*::GFP worms exposed to each bacterial treatment (e) Representative fluorescent images showing worms exhibiting cytoplasmic, intermediate, and nuclear DAF-16 localization. (f) Heatmap displaying standardized Pearson residuals calculated using the contingency table. Positive residuals indicate more worms in a localization category than expected and negative residuals indicate fewer than expected. Asterisks indicate significant deviations from expected frequencies, with \**P* < 0.05, \*\**P* < 0.01, and \*\*\**P* < 0.001. (g) A global Pearson chi-square test was used to assess whether DAF-16 localization differed among bacterial treatments (*P* = 0.0001). (h) Fluorescent images of *C. elegans* carrying *gst-4p*::GFP exposed to *E. coli*, *Alishwanella sp.nov*., *C. flavum*, or *E. roggenkampii*. (i) Quantified mean fluorescence intensity of *gst-4p*::GFP worms exposed to each bacterial treatment. Statistical significance in *b, d, i* was determined by one-way ANOVA followed by Dunnett’s test comparing each strain with the *E. coli* OP50 control.

### Wastewater isolates activate host defense pathways

To assess the host response to potentially pathogenic wastewater isolates, a PMK-1/p38 MAPK activation reporter strain was used (Fig. 5c). The PMK-1/p38 MAPK pathway is one of the most frequently activated host innate immune response pathways in *C. elegans*, particularly during bacterial colonization of the intestine^17^. Expression of the PMK-1-dependent reporter, *irg-5p*::GFP, varied among isolates, indicating differences in host immunity activation, with *P. aeruginosa, Alishewanella sp.nov., C. amalonaticus, E. roggenkampii, A. towneri* and *S. xiamenensis* all exhibiting enhanced fluorescence (Fig. 5d). Interestingly, decreases in median lethal time (LT50) did not completely correspond with increased innate immune activation (compare 5b to 5d). This may indicate that some of these isolates evade or suppress canonical innate immune responses, or reflects the initiation of disease processes that are poorly monitored by the PMK-1 pathway.

We also investigated the potential of these isolates to trigger activation of host stress-response signaling pathways, a common consequence of intestinal infection in *C. elegans*. To test this, a DAF-16::GFP strain was used. DAF-16/FOXO is a transcription factor whose nuclear localization indicates the activation of insulin signaling, and is associated with strong resistance to multiple biotic and abiotic stressors^18^. Individual worms were scored based on cytoplasmic, intermediate, or nuclear DAF-16 localization (Fig. 5e). Although significant differences in localization patterns were observed, only *P. guangdongensis* triggered a strong translocation of DAF-16 to the nucleus (Fig. 5 f-g). Several other isolates displayed increased cytoplasmic localization, suggesting inhibition of insulin signaling under these conditions. To assess the effect of these isolates on host oxidative stress and detoxification responses, a *gst-4*::GFP strain was used^19^ (Fig. 5h). Our results showed that *Alishwanella sp.nov.*, *C. flavum* and *E. roggenkampii* increase *gst-4p*::GFP expression, indicating an induction of oxidative stress or activation of detoxification responses. In contrast, other isolates produced minimal activation despite reducing host survival, suggesting alternative modes of host defense or immune evasion (Fig. 5i). Overall, our results demonstrate that wastewater-derived isolates infect *C. elegans* and activate host immune and defense systems through diverse mechanisms. Across the nine isolates, pathogenicity could not be predicted by activation of a single immune pathway.

### Human epithelial cell adhesion was uncommon; antimicrobial resistance was widespread

We also screened the nine isolates for colonization potential and antimicrobial resistance (Fig. 6; Table S10, Table S11). Only two adhered detectably to the human epithelial cell line A549: *C. kerstersii* at 8.3 ± 2.6% of the inoculum and *P. guangdongensis* at 7.3 ± 4.7%, against 11.7 ± 9.4% for the *P. aeruginosa* PA01 positive control and 0.6 ± 0.6% for the *E. coli* C600 negative control (one-sided Mann-Whitney, Benjamini-Hochberg adjusted *P* = 0.039 for both). Antimicrobial susceptibility testing against eight antibiotics spanning six mechanistic classes showed that four isolates, *C. kerstersii*, *C. flavum*, *L. capsici* and *S. xiamenensis*, were resistant to at least half the panel. *C. flavum* was resistant to all eight, consistent with the intrinsic resistance previously reported for this genus²¹.

**Fig. 6.**
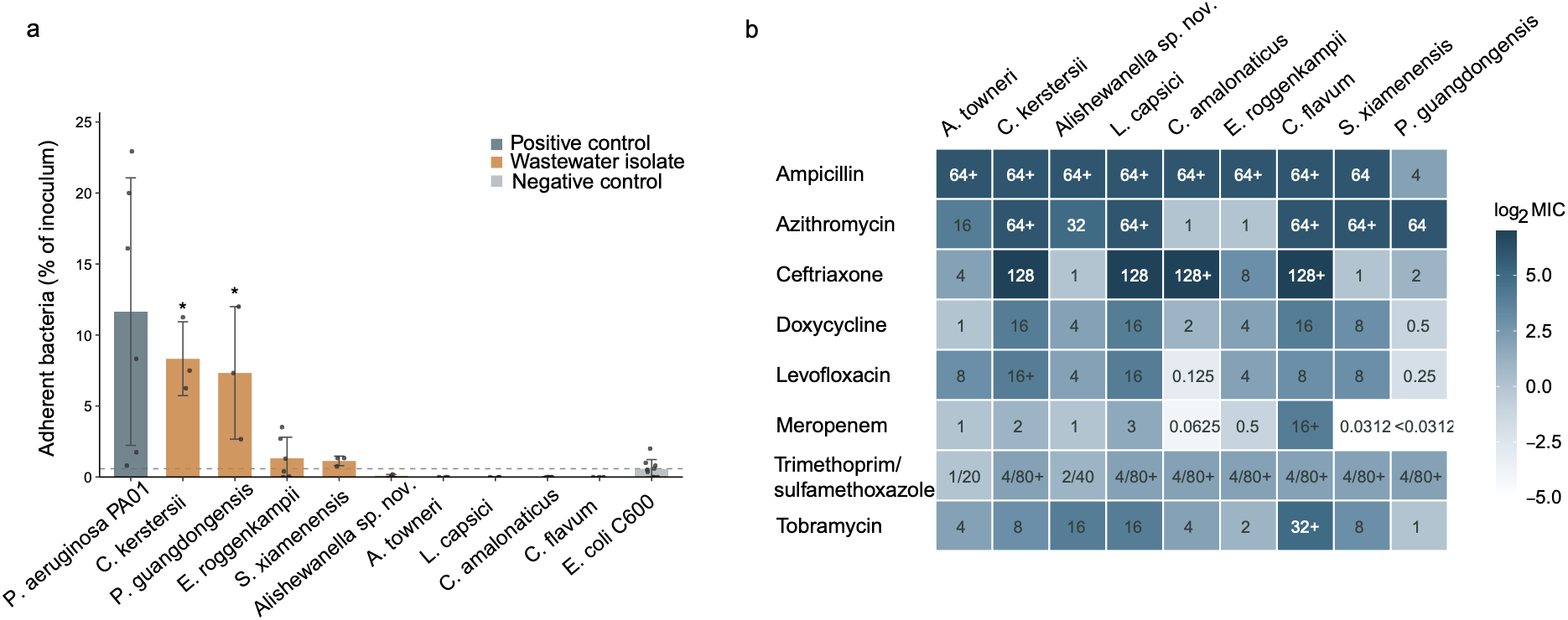
Adhesion and antibiotic susceptibility of the nine isolates. (a) Adhesion to A549 epithelial cells. Adherent bacteria as a percentage of the inoculum after incubation with A549 monolayers. Bars give the mean, error bars the standard deviation, and points the individual replicate wells; replicate numbers differ between strains because wells with off-target multiplicity of infection were excluded. The dashed line marks the mean of the *E. coli* C600 negative control. Asterisks mark isolates adhering significantly above that control (one-sided Mann-Whitney, Benjamini-Hochberg adjusted *P* < 0.05). Wells in which no adherent colonies were recovered are plotted as zero and represent values below the limit of detection. (b) Minimum inhibitory concentrations for eight antibiotics spanning six mechanistic classes, determined by broth microdilution in cation-adjusted Mueller-Hinton medium following a modified CLSI M07 protocol, in triplicate. Cell color encodes log₂ MIC and the printed value is the MIC in µg/mL. A trailing plus sign indicates that the isolate grew at the highest concentration tested, so the MIC is a lower bound rather than a measured value; this applies to 23 of the 72 measurements, including seven of nine for ampicillin and seven of nine for trimethoprim-sulfamethoxazole, neither of which therefore discriminates among these isolates. Trimethoprim-sulfamethoxazole was tested as a 1:20 combination and is reported as both components. Values for both panels are provided in Supplementary Table S10 and Supplementary Table S11.

## Discussion

Molecular wastewater surveillance has become an essential tool for tracking known pathogens, antimicrobial resistance genes and sequence-defined variants, but its sensitivity to emerging threats is constrained by the targets and reference databases used for detection^20–22^. PCR-based assays require predefined markers, and sequence-based approaches prioritize organisms or genes that resemble previously characterized pathogens^8^. This creates a blind spot for viable PEPs that are rare, poorly represented in reference databases, or cause damage via mechanisms that are not yet annotated^6,23^. Our study addresses this limitation by pairing wastewater surveillance with an untargeted functional enrichment step that selects for growth in human serum .

Using human serum for selection, Capture-Enrich successfully captured a wide-range of known pathogens, commensals, and PEPs from community wastewater. Serum passaging restructured wastewater communities and recovered ASVs that were not detected by direct amplicon sequencing of raw wastewater, indicating that functional enrichment can reveal rare serum-prolific lineages that may be missed by sequencing alone. Capture-Enrich adds a complementary culture-based functional layer that can prioritize organisms for deeper characterization.

Capture-Enrich further illustrates how marker-based surveillance may under-sample environmentally circulating lineages with clinical relevance. A majority of our serum-selected isolates consisted of novel sequence types (STs) or belong to genera lacking standard MLST surveillance frameworks, potentially placing them outside routine sequence-type tracking. Canonical clinical marker tools failed to prioritize highly adjacent *K. pneumoniae* environmental isolates due to a lack of traditional hypervirulence or high-risk carbapenem-resistance signatures. Integrating functional screens such as Capture-Enrich into surveillance infrastructure would help close those gaps, both by characterizing immune-evasion phenotypes directly and by supplying new sequences of concern to the reference sets that sequence-based tracking depends on.

Wastewater-derived ESKAPE isolates were interleaved directly with clinical reference genomes rather than forming distinct, isolated environmental clades. The identification of concerning lineages sharing clinical sequence types, with nearest-neighbor distances of 157–310 SNPs across the shared core genome, indicates close but non-identical relatedness between some wastewater and clinical lineages. This structural continuum supports a model wherein ostensibly non-pathogenic microbes could adapt to colonization of humans as opportunistic pathogens or that these strains are already pathogenic but have not yet emerged. Unsurprisingly, many of the PEPs we isolated, such as *C. aquatica*^24^, *C. amalonaticus*^25–27^ and *S. xianmensis*^28^ have very few patient cases, though it is also likely that such rare infections may not be identified correctly. Typically, these patients are immunocompromised. Immunodeficiency can provide a weakened selection environment for PEPs, allowing them to adapt to the host and effectively "climb the ladder" toward systemic pathogenicity within human hosts.

We characterized 161 isolates to identify phenotypic and genomic features associated with robust serum growth, which we used as a functional proxy for evasion of humoral innate immunity. Thirteen isolates grew strongly in serum despite minimal damage activity, and 55 isolates grew in serum without carrying any curated complement-evasion or serum-resistance gene. The discordance was not specific to serum: bile-salt tolerance was present in 129 isolates and was likewise unexplained by any recognizable hydrolase gene. This systemic disconnect highlights an opportunity to expand current database curation for environmental strains that have escaped intensive biomedical scrutiny, making a compelling case for the further investigation of PEPs.

A deeper analysis of nine environmental isolates confirmed that our functional serum screen reliably identifies lineages with active pathogenic potential. All selected PEP candidates significantly shortened host lifespan in *C. elegans* infection models. However, accelerated mortality did not always map neatly to standard host immune activation. This distinction suggests that these environmental strains may deploy unique, uncharacterized strategies to evade, or actively suppress, canonical host innate defenses. Environmental strains actively triggered host stress responses and selectively altered responses in metabolic signaling pathways that closely mirrored the virulent control strain. *P. guangdongensis* and *C. kerstersii* showed moderate epithelial cell adherence. We speculate that adherence is a bespoke adaptation to the host and would thus be less common among PEPs that have not adapted to a specific host range^29^. These candidate PEPs predominantly match the profiles of opportunistic pathogens. They are functionally equipped to capitalize on compromised or weakened host immunity, to potentially establish early-stage infection processes. Ultimately, integrating functional screening platforms like Capture-Enrich alongside established genomic frameworks could enhance the depth of environmental biosurveillance, offering a scalable and agnostic strategy to functionally validate uncurated phenotypes and proactively enrich the reference databases that sequence-based tracking relies upon.

## Methods

### Wastewater collection and serum exposure

Raw wastewater was collected from two municipal wastewater treatment plants (WWTPs) in Houston, Texas, USA, designated Site A and Site B. Site A is a large advanced-secondary plant that uses a two-stage, pure-oxygen activated-sludge process, with a permitted daily average flow of approximately 200 million gallons per day (MGD), and serves a mixed catchment encompassing the central business district together with commercial and industrial areas. Site B is an activated-sludge plant serving a predominantly residential catchment. Samples were collected as 24-h composite samples of raw influent wastewater in February 2024 for the serial passages over 18 days. For each site, 100 ml of wastewater was collected in autoclaved polypropylene bottles, transported to the laboratory on ice, and processed within 2 h of collection.

Wastewater was pre-filtered through a PTFE membrane filter (Sigma-Aldrich, 10 µm pore size) to remove larger particles, and cells were pelleted by centrifugation. Pelleted cells were resuspended in 100% human serum (GeminiBio) supplemented with 0.8% glucose before encapsulation in 90 µm microdroplets^14,15,30,31^. Glucose was included to support population growth over the three-day incubation. Its effect on population composition was evaluated by metagenomic analysis, which showed little appreciable difference (Fig. S5), although the final OD after three days was consistent with a single additional doubling.

### Microfluidic encapsulation and serial serum enrichment

Flow-focusing microfluidic chips generate encapsulations following a Poisson distribution. To maintain biodiversity, we targeted a mean occupancy (λ) of 0.3 to 1.0 cells per microdroplet, based on prior experiments correlating CFU with optical density at 600 nm (OD600)^14^. Cell suspensions in serum were introduced into the microfluidic chip to generate water-in-oil droplets at an aqueous-phase flow rate of 30 µL/min and an oil-phase flow rate of 100 µL/min. The oil phase consisted of fluorinated oil containing 1.5% (w/w) fluorinated surfactant. The number of microdroplets containing a single cell ranged from approximately 566,800 to 965,800 per mL of emulsion (Table S1). Particulates in the bacterial size range were observed in the initial encapsulation but were removed by dilution in the second iteration. Each microdroplet typically supports 4 to 10 doublings^14,31^. Each population was encapsulated in a 1 mL emulsion.

After encapsulation, emulsions were incubated in a humidified growth chamber at 37 °C for three days. *Bacteroides* were enriched across all six passages. Although they are obligate anaerobes, Capture-Enrich maintained and enriched these populations because the emulsion forms a natural oxygen gradient: aerobes and facultative anaerobes consume oxygen at the upper layers, a phenomenon we have observed previously^30^. Microdroplets were then decapsulated, cells were recovered by centrifugation, and cells were re-encapsulated in fresh serum for the next round of enrichment. After three days of incubation, total population sizes in the decapsulated populations ranged from approximately 3 × 10^7^ to 2 × 10^9^ CFU/mL. Four replicate populations per site were carried through six iterations over 18 days. At each iteration, DNA was prepared for community analysis and population samples were archived at −80 °C for future studies.

### 16S rRNA gene amplicon sequencing and community analysis

Genomic DNA was extracted from raw and serum-passaged wastewater samples using the Maxwell PureFood extraction system (Promega). The V4 hypervariable region of the 16S rRNA gene was amplified using primers 515F/806R with Q5 High-Fidelity DNA Polymerase (New England Biolabs)^32^. Amplicons were purified by gel extraction using the Monarch DNA Gel Extraction Kit (New England Biolabs) and submitted to Azenta Life Sciences for library preparation and paired-end sequencing on the Illumina platform via the Amplicon-EZ service. Each site was sampled once. The wastewater sample was divided into four aliquots; each aliquot was sequenced directly as passage 0 and encapsulated to found one of four parallel enrichment lineages, which were then carried through six serial passages. A total of 62 samples were sequenced, comprising these four lineages at seven timepoints for each of two WWTP sites, plus flask controls at passage 4 (n = 3 per site). Flask controls were excluded from downstream analysis, leaving 56 samples. A further 12 libraries from the glucose-supplementation control were sequenced and analyzed separately (Supplementary Methods).

Demultiplexed reads were processed in QIIME 2 (v 2023.2) using the DADA2 plugin for quality filtering, denoising, merging and chimera removal^33,34^. A mean of 417,501 ± 68,369 raw reads per sample yielded 249,235 ± 44,344 quality-filtered, non-chimeric reads (59.7% retention), producing 16,279 ASVs. Taxonomy was assigned using a naive Bayes classifier (classify-sklearn) trained on the SILVA 138 database (99% OTUs) trimmed to the 515F/806R region^35,36^. A phylogenetic tree was constructed using the align-to-tree-mafft-fasttree pipeline in QIIME 2. Analyses were carried out in R with phyloseq (v 1.46.0) and vegan^37,38^. SILVA left 43% of ASVs without a genus-level assignment, and its literal label "uncultured" spans 105 families in this dataset. Rather than pooling these, each unassigned ASV was labelled by its finest assigned rank, giving 1,184 genus-level bins of which 872 are named genera. Analyses of taxonomic composition and diversity use all 1,184 bins; the screen for genera newly detected during passaging (below) uses named genera only. Library sizes spanned 144,860 to 344,702 reads. Diversity estimates were rarefied to 144,860 reads, the smallest library, so that no sample was excluded. Following Schloss, rarefaction was implemented as the average across many independent subsamples: 1,000 for alpha diversity and the presence-absence analyses, and 100 for Bray-Curtis dissimilarities (vegan::avgdist)^39^. All results were also computed without rarefaction and are compared in Table S14.

Observed ASV richness and genus-level Shannon diversity were compared across passages within each site by Friedman tests blocked on lineage, since the seven timepoints within a lineage are repeated measures on one propagated population. Pairwise comparisons against raw wastewater used Wilcoxon signed-rank tests with Benjamini-Hochberg correction. Beta diversity was assessed on ASV-level relative abundances using Bray-Curtis dissimilarity, visualized by principal-coordinates analysis. Passage and site effects were quantified by PERMANOVA with marginal terms and 9,999 permutations, permutations for passage restricted within lineage^40^. Between-site separation at each passage was quantified with betadisper as the distance between site centroids in principal-coordinate space, expressed relative to the mean distance of lineages to their own centroid^41^.

ASVs were scored as retained from raw wastewater if detected in any of the four raw aliquots from that site, with percentages calculated per lineage then averaged. The three-way overlap between raw wastewater, passage 2 and passage 6 was calculated within each lineage against its own aliquot. A genus was scored as newly detected if absent from all four raw aliquots at that site and present in at least two of four lineages at one or more passages.

### Isolate recovery and whole-genome sequencing

De-encapsulated droplets from iteration 2 of the serum passages were frozen at −80 °C. These samples were thawed, serially diluted and spread-plated. Individual colonies were picked at random from each wastewater site and cultured overnight in brain-heart infusion (BHI) broth. Each culture was streaked onto an individual plate to yield isolated colonies, and single colonies were picked from each plate and cultured overnight in BHI to produce the final collection. Isolates were stored in 25% glycerol at −80 °C and were recovered from freezing by overnight culture in BHI before use in each phenotypic assay.

Recovered isolates underwent whole-genome sequencing on the Oxford Nanopore MinION (FLO-MIN114) with the Rapid Barcoding Kit (SQK-RBK114.96). Reads were assembled with Flye (v 2.9.5-b1801) and polished with Medaka (v1.8.0)^42,43^. Assembly quality was assessed using QUAST (v 5.3.0) and CheckM2 (v1.0.1; completeness ≥ 90%, contamination < 3%), and taxonomic classification was performed with GTDB-Tk (v 2.2.6)^44–46^. Of 185 total assemblies, 161 passed quality control (median 1 contig, N50 4.3 Mb, completeness 99.8%).

### Strain-level typing and clonal group definition

Multilocus sequence typing was performed with mlst (v2.23.0) against PubMLST^47^. Species-specific typing used Kleborate (v 0.3.0; *Klebsiella*), ClermonTyping (*E. coli* phylogroups) and ECTyper (v 2.0.0; *E. coli* serotypes)^16,48,49^. Candidate novel species were defined as isolates with < 95% ANI to the nearest GTDB species.

Clonal groups were defined in two steps. First, all-versus-all ANI was computed with FastANI (v1.33); pairs sharing ≥ 99.9% ANI and aligned fraction (AF) ≥ 0.90 were grouped by single-linkage; every qualifying pair exceeded the AF cutoff (minimum 0.946). Each candidate cluster was then validated by within-species core-genome SNP analysis (0.95 core threshold, snp-dists v1.2.0); members separated by more than 100 core SNPs were split apart or reclassified as singletons^50,51^. This assigned 71 of the 161 isolates to 21 clonal groups of two to eight members, leaving 90 singletons and 111 distinct genotypes overall.

### Phylogenomic comparison with clinical reference genomes

To assess clinical relevance, wastewater isolates from the four most abundant ESKAPE-adjacent species, *P. aeruginosa* (n = 32), *K. pneumoniae* (n = 7), *E. coli* (n = 5) and *E. hormaechei* (n = 4), were compared with all complete-assembly reference genomes available for each species in the CDC HAI-Seq surveillance collection (BioProject PRJNA288601; accessed January 2025; 59, 48, 50 and 22 references, respectively). Wastewater and reference genomes were annotated with Prokka (v 1.14.6), per-species core-gene alignments were built with Panaroo (v 1.5.2; strict mode, 0.95 core threshold), and maximum-likelihood phylogenies were inferred with RAxML (v 8.2.12) under the GTRGAMMA model^52–54^. Trees were visualized in iTOL^55^.

### Functional annotation with SeqScreen and FunSoC

Genome assemblies from the 161 isolates were screened for Functions of Sequences of Concern (FunSoC) using SeqScreen with SeqScreenDB (v 23.4)^56^. SeqScreen was run separately for each isolate with windowed screening enabled. SeqScreen annotations were filtered to retain high-confidence calls, defined as hits with a best e-value ≤ 1 × 10^−5^. For each isolate, retained gene calls were summarized across the 32 FunSoC categories to generate an isolate-by-category count matrix. A FunSoC category was considered present in an isolate if at least one high-confidence gene assigned to that category was detected.

### Phenotypic assays

Isolates were assayed for a panel of *in vitro* phenotypes associated with host interaction, comprising four secreted-damage traits (hemolysis, gelatinase, caseinase and DNase activity) and two persistence traits (bile-salt tolerance and biofilm formation). Growth in human serum was assessed separately.

### Hemolysis assay

Samples were streaked from overnight culture onto blood agar base (Thermo Scientific) containing 6% (v/v) sheep’s blood, incubated overnight at 37 °C with 5% CO_2_, and scored for alpha, beta or gamma hemolysis. All assays were performed in triplicate.

#### Gelatinase Assay

The gelatinase assay was performed with modifications^57^. Samples were spotted onto Todd-Hewitt agar plates (Sigma-Aldrich) containing 3% (w/v) gelatin from bovine skin using a 96-well replicator, incubated at 37 °C, developed with 3.5 M ammonium sulfate, and observed for a halo around the sample. All assays were performed in triplicate.

#### Caseinase Assay

As for the gelatinase and DNAse assays, samples were spotted onto a 15 cm agar plate containing 28 g/L skim milk powder, 5 g/L tryptone, 2.4 g/L yeast extract, 1.0 g/L glucose and 15 g/L agar using a 96-pin replicator. Plates were incubated overnight at 37 °C and colonies were observed for a clear halo, indicating caseinase production. All assays were performed in triplicate.

#### DNAse Assay

Isolates were spotted onto a 15 cm dish of Difco DNase Test Agar (Fisher Scientific) using a 96-pin replicator. Plates were developed with 1 M HCl, and samples were observed for a clear halo, indicating DNase production. *Pseudomonas aeruginosa* developed a divergent phenotype, with colonies turning pale pink and generating no halo; *P. aeruginosa* was therefore recorded as negative in this assay, although prior studies have documented *P. aeruginosa* nuclease production^58^. All assays were performed in triplicate.

#### Bile salt tolerance assay

The bile salt survival assay was performed with modifications^59^. Samples were grown to log phase and diluted 1:100 into brain-heart infusion (BHI) broth, BHI with 0.125% (w/v) bile salts, and BHI with 0.5% (w/v) bile salts. Cultures were grown for 12 h at 37 °C with 5% CO_2_, and OD600 was measured in an Epoch 2 microplate reader. OD600 in 0.5% (w/v) bile salt was compared with OD600 in BHI for each sample. All assays were performed in triplicate.

#### Biofilm Assay

The biofilm assay was performed in BHI (Difco) as described^60^. Samples were cultured in a non-tissue-culture-treated plate for 24 h at 37 °C, rinsed, stained with crystal violet for 10 min, and rinsed again. Once dry, wells were qualitatively scored for the appearance of a stained biofilm ring. All assays were performed in triplicate.

#### Serum Growth Assay

A total of 161 isolates were evaluated for their ability to grow in 100% human serum (GeminiBio). Isolates were inoculated into individual wells of 96-well plates containing 200 µL of 100% human serum per well. Plates were incubated overnight at 37°C with orbital shaking at 100 rpm. Growth was determined by measuring optical density at 600 nm (Epoch 2 microplate reader) with initial baseline values subtracted from final readings. All assays were performed in triplicate.

### *C. elegans* strains and maintenance

All *C. elegans* strains were maintained on standard nematode growth medium (NGM) seeded with *E. coli* (OP50) and maintained at 22°C^61^, unless otherwise noted. *C. elegans* strains used in this study include SS104 [*glp-4*(*bn2*)]^62^, AY101 |*acIs101* [*pDB09.1*(*irg-5p*::GFP); *pRF4*[*rol-6*(*su1006*)]]|^63^, CL2166 |*dvIs19* [(*pAF15*)*gst-4p*::GFP::NLS]|^64^, and NVK265 |*glp-4*(*bn2*); *zIs356* [*daf-16p*::DAF-16a/b::GFP + *rol-6*(*su1006*)]|.

#### Worm synchronization

Worms were synchronized by hypochlorite isolation of eggs from gravid adults, followed by hatching of embryos in S Basal. Approximately 6,000 synchronized L1 larvae were transferred onto 10 cm NGM plates seeded with OP50 and grown at 22 °C for 48 h before experiments, or for three days for propagation. For the SS104 strain, worms were grown at 25 °C for 50 h before experiments, or for five days at 15 °C for propagation.

For all survival assays, young adult *glp-4* worms were used. For assays involving the use of fluorescent reporters, L4-stage worms were used, unless noted otherwise. Bacterial strains used in this study include *P. aeruginosa* PA14^65^, *E. coli* OP50^66^, and multiple wastewater isolates (this study).

#### Slow Killing Assay

Agar-based pathogenesis (Slow Killing or SK) was performed as described^67^. Fifty young-adult worms were transferred onto bacterial lawns on SK plates and incubated at 25 °C. Dead worms were scored daily to generate survival curves, and worms found outside the agar were censored. *E. coli* OP50 was used as a non-pathogenic negative control, and *P. aeruginosa* PA14 was used as a positive control. At least three biological replicates were performed for all assays.

#### Fluorescent reporter imaging and quantification

Exposure of fluorescent-reporter worms to bacterial isolates was performed as for the slow-killing assay. At the L4 stage, 500 worms were placed onto bacterial lawns for 24 h. Worms were then washed off the plates and sorted into 96-well plates or onto 3% (w/v) Noble agar slides for imaging. Worms were paralyzed using 10 mM levamisole hydrochloride. At least three biological replicates were performed for all assays.

For reporter strains AY101 and CL2166, 50 worms per well were sorted into 96-well plates and imaged using a Cytation 5 Cell Imaging Multi-Mode Reader (BioTek Instruments). Mean GFP expression was calculated per well. At least three biological replicates were performed for all assays.

For visualization of *glp-4(bn2)*;DAF-16::GFP, worms were immobilized with 10 mM levamisole and mounted on 3% (w/v) Noble agar slides. At least 15 worms per replicate were imaged, with three biological replicates. All images were captured on a Zeiss ApoTome.2 Imager.M2 (Carl Zeiss) at 20x magnification. Images were scored manually and classified as nuclear, intermediate or cytoplasmic DAF-16 localization.

### Mammalian cell culture and adhesion assay

#### Cell culture

A549 cells (ATCC CRM-CCL 185) were cultured in high-glucose Dulbecco’s Modified Eagle Medium (DMEM) (Gibco) containing 10% (v/V) Fetal Calf Serum (FCS) (Gibco) and 1% penicillin-streptomycin at 37 °C with 5% CO_2_. Cells were passaged every 5 to 7 days at 80 to 90% confluence for at least three passages after thawing. For the adhesion assay, cells were seeded at 1.5×10^5^ cells/mL in a 24-well plate in high-glucose DMEM containing 10% FCS and no antibiotics, and grown for 48 h to a confluent monolayer.

#### Adhesion assay

Cells were used for the assay between passages 15 and 25. The assay was performed with modifications^68^. Once cells reached a confluent monolayer, DMEM containing 10% (v/v) FCS was replaced with high-glucose DMEM containing no FCS. Cells were inoculated with bacterial samples at a Multiplicity of infection (MOI) of 25 to 40. A well containing 1 mL DMEM and no cells received the same inoculum as a reference. The cell and bacterial suspension were incubated for 1 h at 37 °C with 5% CO_2_. Cells were then washed three times with Dulbecco’s phosphate-buffered saline (DPBS) and lysed for 15 min with 1% (v/v) Triton X-100 in DPBS. Lysates and the cell-free inoculum reference were serially diluted and plated for CFU enumeration. Adhesion was reported as the percentage of the inoculum recovered in the lysate. All measurements were performed in at least triplicate using *P. aeruginosa* PAO1 (positive) and *E. coli* EC600 (negative) as controls.

### Antibiotic susceptibility testing

Minimum inhibitory concentrations (MICs) were determined for eight antibiotics: ampicillin, azithromycin, ceftriaxone, doxycycline, meropenem, levofloxacin, trimethoprim/sulfamethoxazole, and tobramycin (Sigma-Aldrich). Testing followed a modified version of the CLSI M07 method for dilution antimicrobial susceptibility tests for bacteria that grow aerobically^69^. An overnight culture of each strain was diluted to OD 0.01 and then 1:200 into fresh cation-adjusted Mueller-Hinton medium (Sigma-Aldrich). 140 µL of this culture was deposited into 96-well plates in triplicate. Antibiotics were added and drug gradients prepared by serial dilution using a Hamilton Microlab Prep liquid-handling robot to a final volume of 150 µL per well. Plates were incubated overnight at 37 °C, after which optical density was measured and background subtracted. The MIC was taken as the lowest drug concentration at which there was no visible turbidity, corresponding to an OD below 0.05. Concentration ranges were set from the predicted ranges of efficacy given in CLSI M100^70^. *E. coli* ATCC 25922, *P. aeruginosa* ATCC 27853, *S. aureus* ATCC 29213, and *E. faecalis* ATCC 29212 were included on each plate as quality-control strains, and their MICs were confirmed to fall within the CLSI-specified ranges before the corresponding test results were accepted. Clinical breakpoints are not established for these environmental species, so susceptibility was interpreted relative to two laboratory reference strains rather than against defined thresholds. *P. aeruginosa* PA14 was used as the reference for meropenem, ceftriaxone, levofloxacin, and tobramycin, and *E. coli* OP50 for ampicillin, azithromycin, doxycycline, and trimethoprim/sulfamethoxazole.

## Statistical analysis

### Damage Phenotype Index (DPI)

Each isolate was characterized across six phenotypic assays grouped into two functional classes. The secreted-damage class comprised hemolysis, gelatinase, caseinase and DNase activity. The persistence class comprised bile-salt tolerance and biofilm formation. Growth in human serum was recorded separately as the enrichment-selection phenotype and was not included in the DPI. Each assay was scored on an ordinal scale (0, 1, 2) and binarized to presence (score ≥ 1) or absence (score = 0). The DPI was calculated as a weighted sum of the six binarized calls, with secreted-damage assays weighted twice relative to persistence assays:

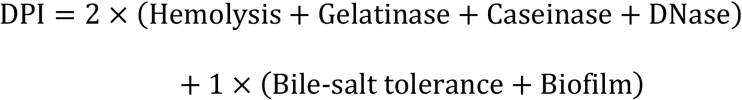

DPI values ranged from 0 to 10, with the secreted-damage component contributing 0 to 8 and the persistence component contributing 0 to 2.

### Phenotype distribution and structure

All phenotype analyses were performed in R (v4.3.1) on the working cohort of 161 isolates. Growth in human serum was measured as OD600 and scored as positive at OD600 ≥ 0.3. The association between DPI and serum growth was assessed by Spearman rank correlation. Pairwise associations among the seven phenotypes were quantified by Spearman rank correlation, with *P* values adjusted across pairwise comparisons using the Benjamini–Hochberg false-discovery-rate procedure^71^. Phenotypic structure was summarized by principal-component analysis on the seven phenotypes, centered and scaled to unit variance, with PC1 oriented to increase with DPI. The contributions of species and sampling site to phenotypic variation were tested by permutational multivariate analysis of variance (PERMANOVA, adonis2; vegan, v2.7-1) on Euclidean distances computed from the six binarized assays and z-scored serum growth, with species and site evaluated as marginal terms over 999 permutations under a fixed random seed^40^. For the species-level heatmap, mean profiles were computed for species represented by at least five isolates and ordered by hierarchical clustering of Euclidean distances with average linkage. Isolates showing strong serum growth with low damage potential were defined as those with serum OD600 ≥ 0.3 and DPI ≤ 2.

#### Genotype-phenotype concordance

For each assay, the phenotype call was compared with the presence of matching annotated genes in the genome. Concordance was tested with Fisher’s exact tests on 2 × 2 tables of phenotype call versus gene presence, and odds ratios and Youden’s *J* index (sensitivity + specificity − 1, treating gene presence as the predictor of phenotype positivity) were reported for each assay^72^.

#### Nematode survival and reporter assays

Slow-killing survival data were plotted as survival curves and analyzed in R (v 3.6.3). Differences between survival distributions were assessed by the log-rank (Mantel-Cox) test with Bonferroni-adjusted pairwise comparisons. LT50 values were calculated with a custom R script based on linear interpolation of the survival data. Differences in GFP reporter activation among bacterial treatments were assessed by one-way analysis of variance (ANOVA) followed by Dunnett’s post hoc test.

DAF-16 localization distributions were compared across treatments with a global Pearson chi-square test of independence. When expected cell counts were small, p-values were estimated by Monte Carlo simulation with 10,000 replicates. The global test was followed by standardized-residual analysis to identify localization categories contributing to the overall association. Positive residuals indicate enrichment of a category relative to the null expectation of independence, and negative residuals indicate depletion. The significance of individual residuals was estimated from the standard normal distribution and adjusted for multiple testing with the Benjamini-Hochberg false-discovery-rate correction^71^. Standardized residuals and their significance levels are displayed in the heatmap. Throughout, statistical significance is indicated as: ns, *p* > 0.05; *, *p* < 0.05; **, *p* < 0.01; ***, *p* < 0.001.

## ACKNOWLEDGEMENTS

This study was supported by NIH/NIAID grant R21AI190686 to YS and NIH/NIAID grant R21AI190938 to LS, YS, and TT. We thank the Houston Health Department and Houston Water for providing wastewater samples.

## DATA AVAILABILITY

The 16S rRNA gene amplicon sequencing and whole-genome sequencing data generated in this study have been deposited in the NCBI under BioProject accession PRJNA1476809.

