## Supplementary material for "Microfluidic Capture-Enrichment of individualized microbes from urban wastewater reveals a hidden reservoir of potential pre-emergent pathogens": Fig. S1; Fig. S2; Fig. S3; Fig. S4; Fig. S5; Supplementary Methods; Table S1

**Affiliations:**

### Supplementary Figures

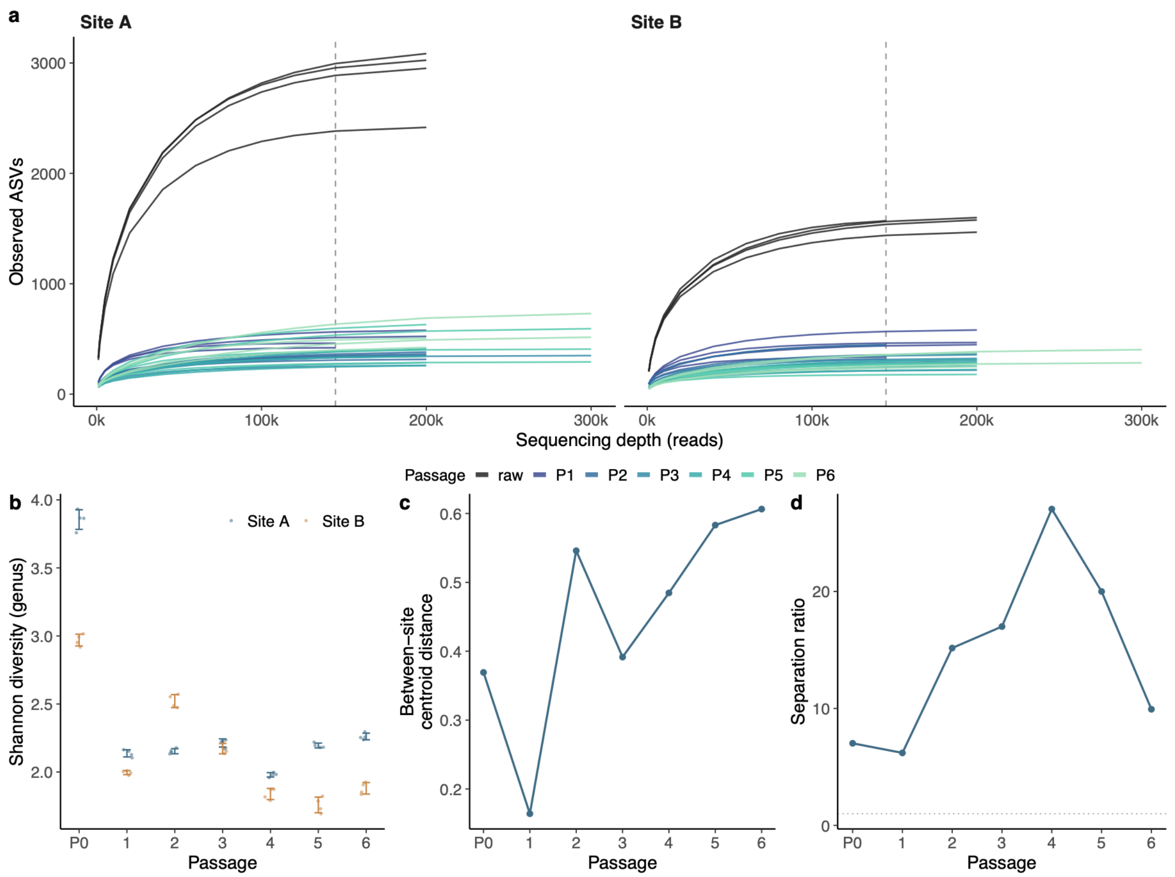


**Fig. S1 Sequencing depth, evenness and between-site separation across serum passages.** All panels use the 56 amplicon libraries from raw wastewater (P0) and six serial serum passages (P1 to P6) at two WWTP sites. Each site was sampled once and the sample divided into four aliquots, each founding one parallel enrichment lineage, so the four values at each passage are lineages rather than independent samples. (a) Rarefaction curves for each library, faceted by site. Curves are colored by passage, with raw wastewater in grey. The dashed line marks 144,860 reads, the smallest library size, to which all diversity estimates were rarefied. (b) Shannon diversity at genus level. Points are individual lineages, bars give mean ± s.d.. Diversity was calculated on natural logarithms after collapsing to genus, retaining ASVs unresolved at genus level as separate bins labelled by their finest assigned rank, and averaged across 1,000 independent subsamples. (c) Distance between the Site A and Site B community centroids at each passage, computed with betadisper as the Euclidean separation in the principal-coordinate space derived from Bray-Curtis dissimilarities of ASV-level relative abundance. Separation was lowest after the first passage (0.164) and highest at passage 6 (0.606), and the trajectory between these points was not monotonic. (d) The same distance divided by the mean distance of lineages to their own site centroid. The dotted line marks a ratio of one, at which the two sites would differ by no more than lineages of a single site do. The ratio remained above 6 at every passage and peaked at 26.4 at passage 4.

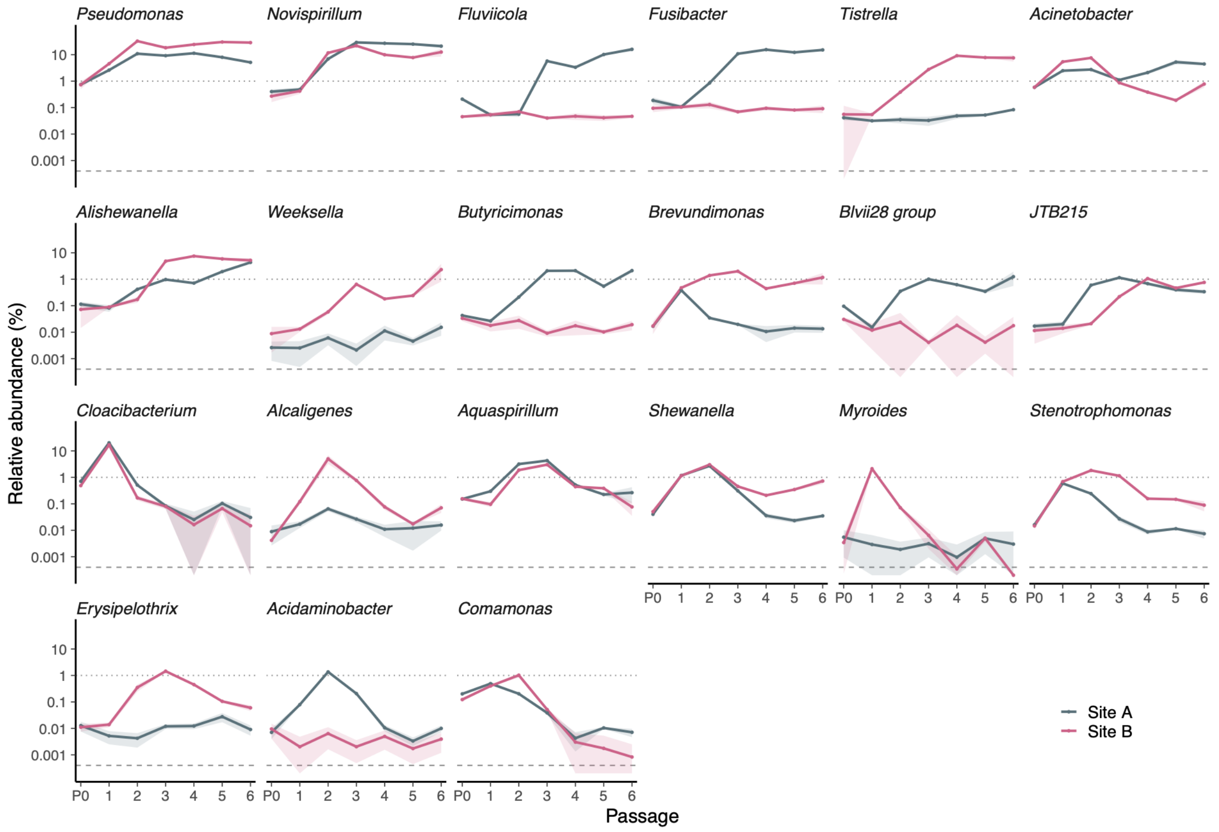


**Fig. S2 Serum passaging expands lineages that were rare in raw wastewater.** Relative abundance across raw wastewater (P0) and six serial serum passages at two WWTP sites, shown for the 21 genera that were below 1% in raw wastewater at both sites and exceeded 1% at some passage at either site. Panels are ordered by peak abundance. Lines give the mean of four biological replicates and shaded ribbons the standard deviation; the y axis is logarithmic. The dotted line marks 1% of the community and the dashed line the relative abundance of a single read at the mean library size (0.0004%), below which a genus could not be detected. Expansions spanned 5-fold to 562-fold. *Alcaligenes* rose from 0.009% to 5.0% and fell back to 0.07%, *Weeksella* from 0.009% to 2.3%, and *Tistrella* from 0.056% to 9.1%. *Cloacibacterium* reached 20.2% at both sites before declining to 0.03% by passage 6. Site-specific responses were common: *Fusibacter* and *Fluviicola* expanded only at Site A, *Tistrella* only at Site B. Two genera carry SILVA placeholder identifiers (*Blvii28_wastewater-sludge_group*, shown as Blvii28 group, and *JTB215*) and have no described species.

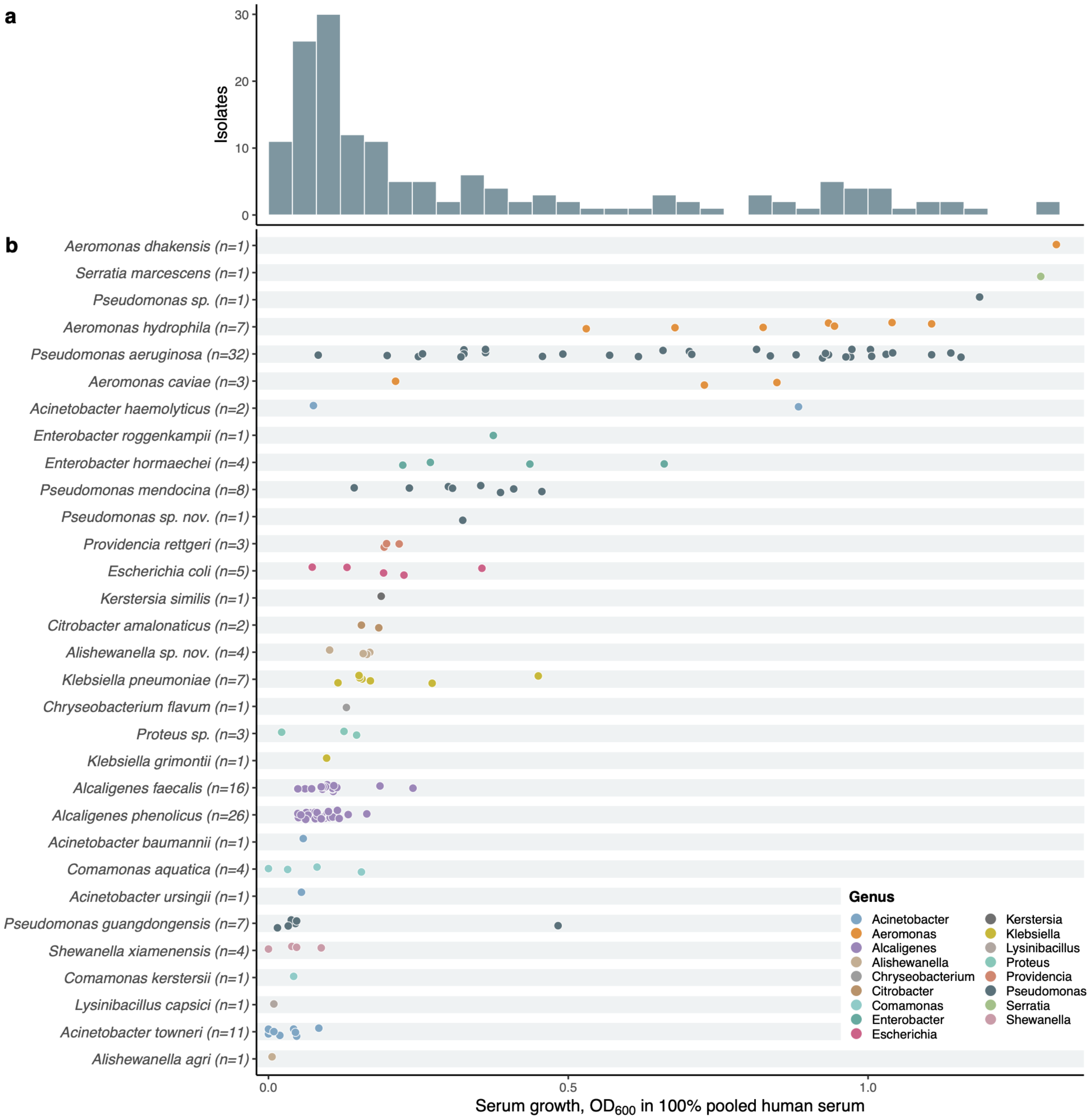


**Fig. S3 Serum growth across the isolate collection.** (a) Distribution of optical density at 600 nm after growth in human serum, all 161 isolates. (b) The same measurements resolved by species, ordered by median. Each point is one isolate, colored by genus; vertical bars mark the species median where n ≥ 3. Species represented by one or two isolates are shown without a median. Values are given in Supplementary Table 8.

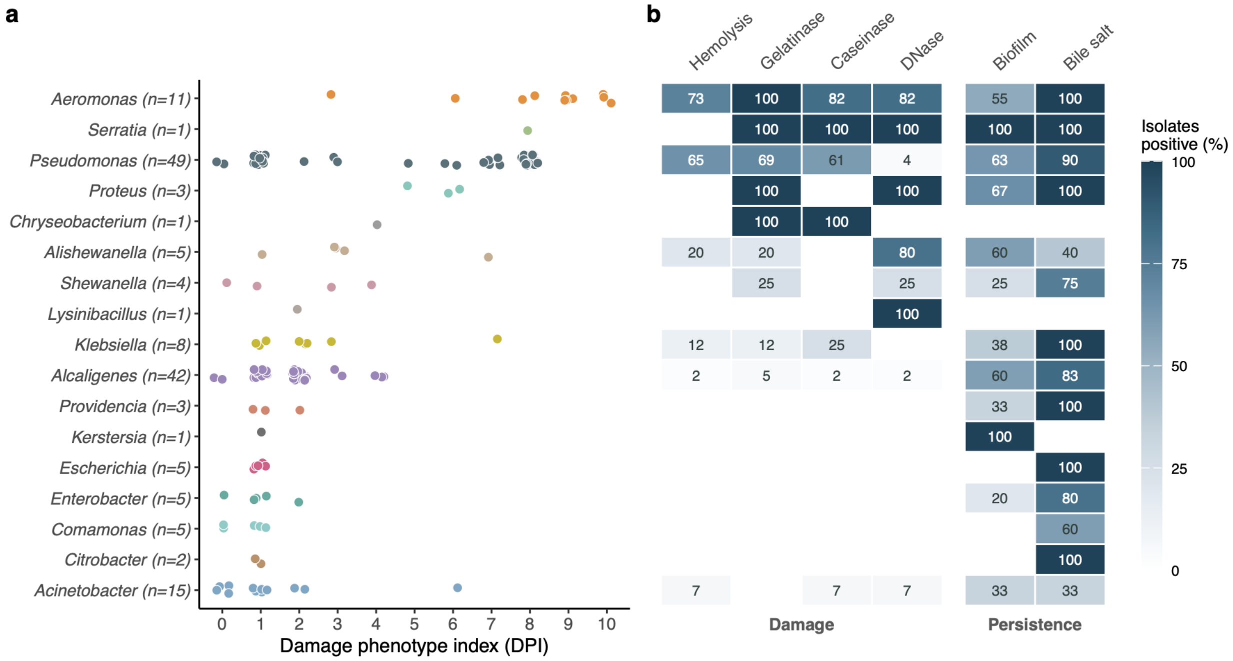


**Fig. S4 Damage phenotype index and assay composition by genus.** (a) DPI for each of the 161 isolates, grouped by genus and ordered by median DPI. Each point is one isolate. (b) Percentage of isolates positive for each assay within each genus, in the same order. Assays are grouped into tissue damage and persistence.

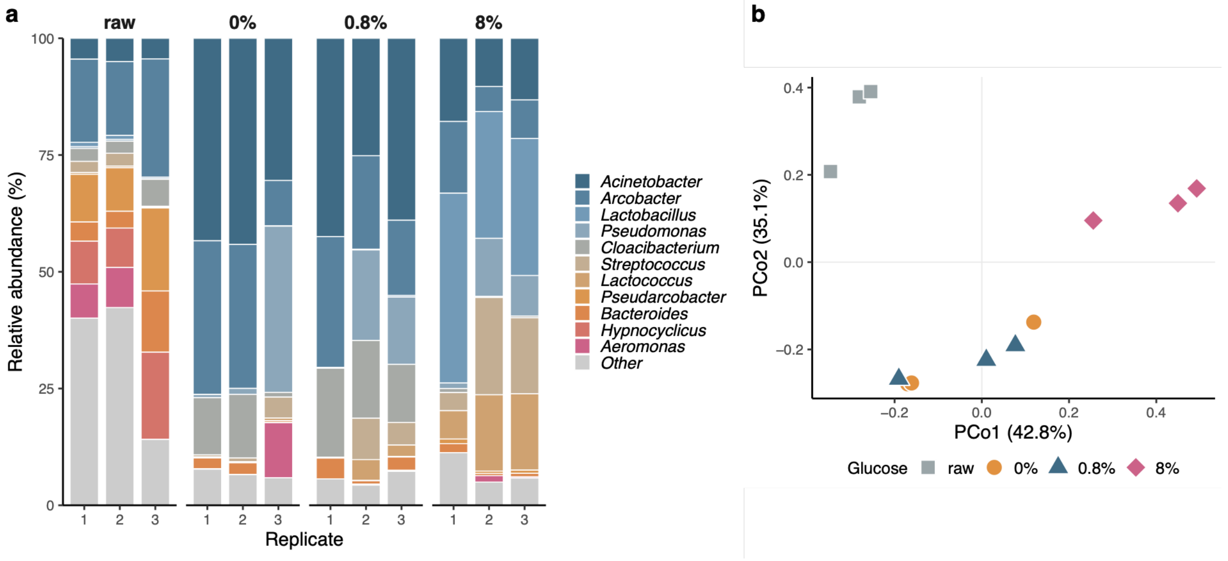


**Fig. S5 Glucose supplementation at 0.8% does not restructure the serum-enriched community.** Site B serum passages at 0%, 0.8% and 8% glucose alongside raw wastewater, three replicate populations per condition, characterized by 16S rRNA gene amplicon sequencing. (a) Genus-level composition of each replicate; the eleven most abundant genera across all samples are shown individually, all others pooled as Other. (b) Principal coordinates analysis of Bray-Curtis dissimilarity between genus-level profiles. Composition at 0% and 0.8% glucose was closely similar across the abundant taxa (*Acinetobacter* 39.3% versus 35.5%, *Arcobacter* 24.5% versus 21.4%, *Pseudomonas* 12.5% versus 11.3%, *Cloacibacterium* 9.0% versus 16.0%), and the two conditions overlap in b. At 8% glucose the community was taken over by fermentative lactic acid bacteria: *Lactobacillus* rose from 0.03% at 0% glucose to 32.4%, and with *Streptococcus* and *Lactococcus* accounted for 59.0% of the community. Serum passaging at any concentration reduced genus richness relative to raw wastewater, from 327 genera to between 58 and 78. Dissimilarity statistics are given in Supplementary Methods.

#### Supplementary Methods

##### Glucose supplementation control

To determine whether glucose supplementation altered which organisms were enriched, a separate passaging experiment was carried out at Site B at three glucose concentrations, 0%, 0.8% and 8% (w/v), with three replicate populations per concentration, alongside three replicates of the corresponding raw wastewater. DNA extraction, amplification, sequencing and denoising followed the procedure described for the main passage series. The twelve libraries yielded 175,352 to 241,106 quality-filtered, non-chimeric reads and 2,937 ASVs. ASVs were collapsed to genus and converted to relative abundance without rarefaction.

Communities were compared by Bray–Curtis dissimilarity on genus-level profiles[1]. With three replicates per condition, formal significance testing has limited ability to support a negative conclusion: for example, a two-sided exact Mann–Whitney comparison of two groups of three has a minimum attainable P value of 0.1, and the corresponding permutation space is similarly limited. We therefore evaluated the magnitude of between-condition dissimilarity relative to the dissimilarity observed among replicate populations within the same condition. Within-condition dissimilarity averaged 0.310 across the three glucose conditions (0.270 at 0.8%, 0.278 at 8%, and 0.381 at 0%). The 0% and 0.8% conditions were separated by a mean dissimilarity of 0.299 across the nine cross-condition pairs, corresponding to 0.96 times the within-condition value. Every other pair of conditions was separated by at least twice that distance (0.62–0.78), including raw wastewater versus each of the three passaged conditions. PERMANOVA showed the same ordering (all four conditions, $R^2 = 0.74$, $P = 0.001$; three glucose concentrations, $R^2 = 0.66$, $P = 0.036$; 0% versus 0.8% alone, $R^2 = 0.08$, $P = 0.76$)[2]. Given the small sample size for the final comparison, we therefore interpreted the magnitude of the Bray–Curtis dissimilarity relative to within-condition variation rather than the nonsignificant P value alone.

#### Supplementary Tables

**Table S1. Single-cell encapsulation efficiency across microdroplet occupancy classes at two cell loading densities.**

| Cells/microdroplet for a 1 ml emulsion | Microdroplets $\lambda = 0.3$ | Microdroplets $\lambda = 1.0$ | Number of cells $\lambda = 0.3$ | Number of cells $\lambda = 1.0$ |
|---|---|---|---|---|
| 0 | 1,944,800 | 985,400 | 0 | 0 |
| 1 | 566,800 | 965,800 | 566,800 | 965,800 |
| 2 | 83,200 | 462,800 | 166,400 | 925,000 |
| 3 or more | 9,400 | 200,000 | 29,000 | 657,800 |
| Total | 2,604,200 | 2,614,000 | 762,200 | 2,548,600 |
